## Supplemental Materials for "Learning fast and fine-grained detection of amyloid neuropathologies from coarse-grained expert labels"

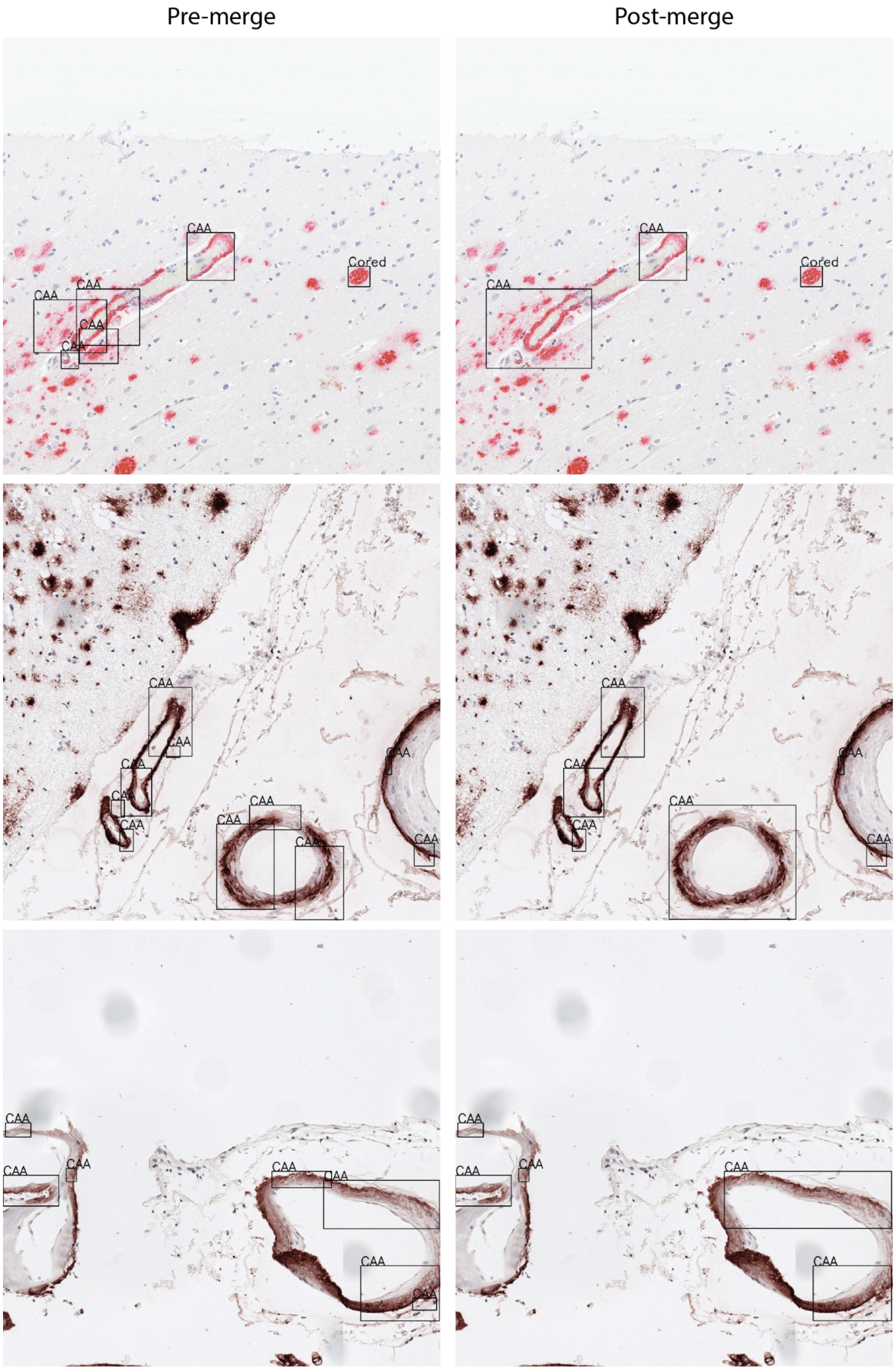


**Supplemental Figure 1: Comparing raw pre-merged labels with merged labels.** Column left shows the raw bounding box labels. Box coordinates were derived from traditional water-shedding methods unassisted by human intelligence. For each identified class shown, at least two out of five expert annotators positively labeled the pathology. Column right shows the merged bounding box labels used to train model version one.

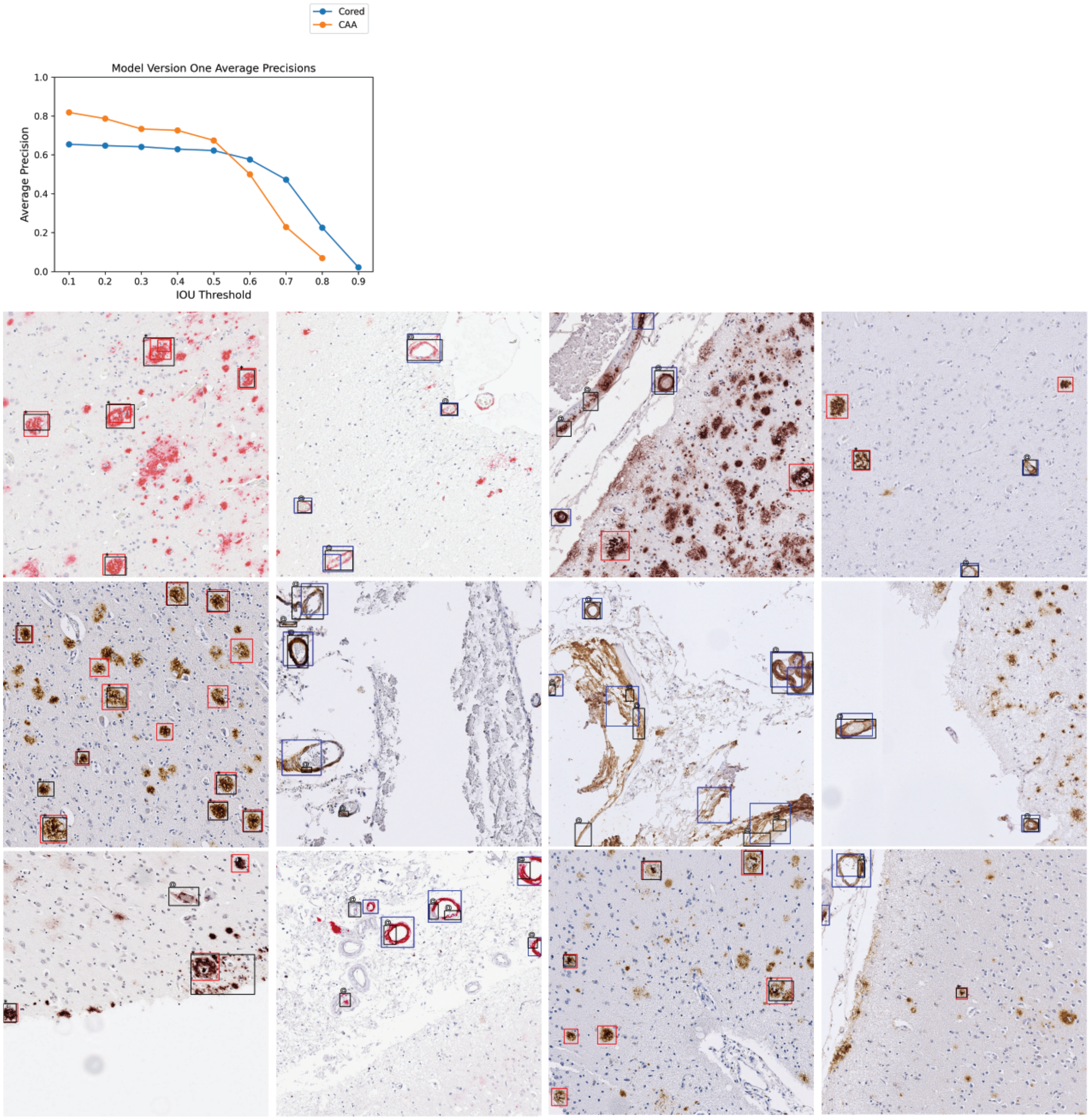


**Supplemental Figure 2: Model version one performance and example image predictions.** Top: Average precisions over the validation set for various IOU thresholds. The AP at IOU=0.90 is undefined for CAA. Bottom: Example images are pulled from the validation set. Cored prediction: red, Cored label: black "*"; CAA prediction: blue, CAA label: black "@". It is important to note that the label data is sparse and does not contain every pathology (Methods).

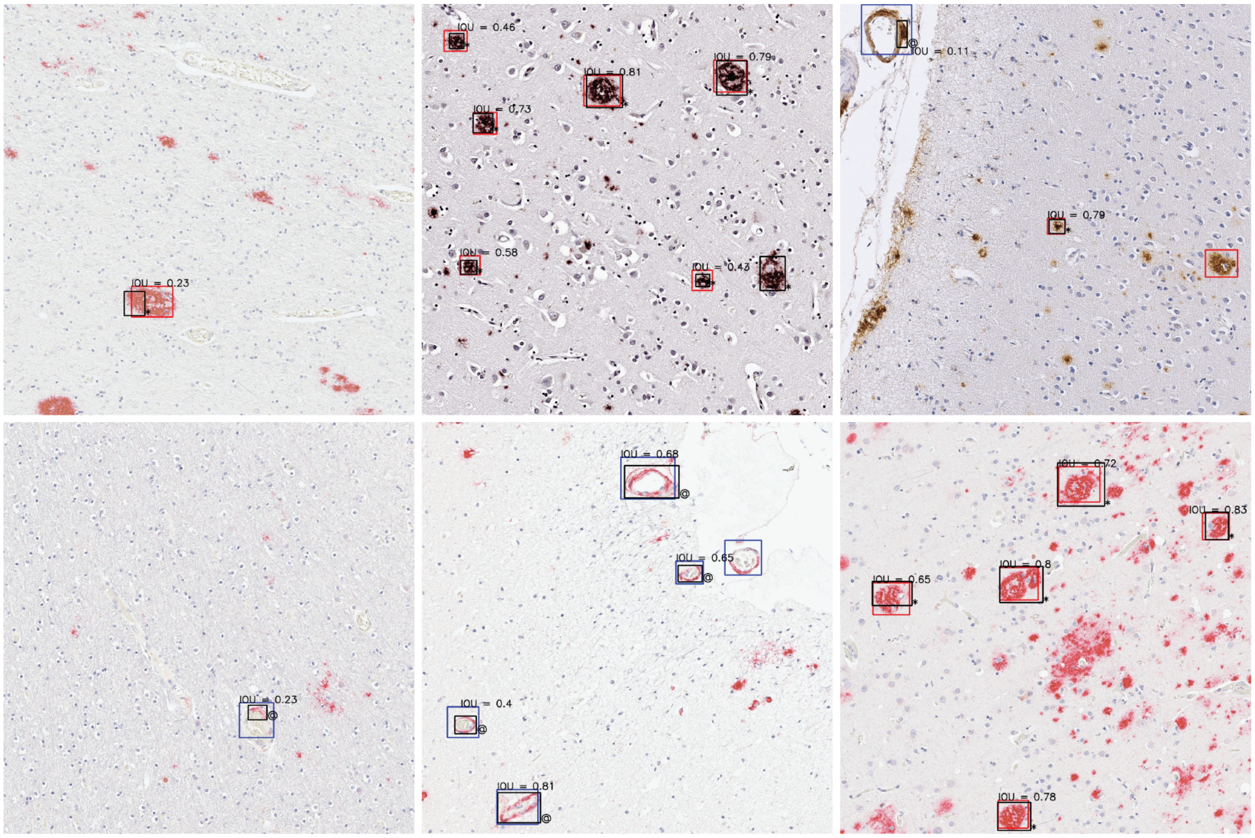


**Supplemental Figure 3: Examples of different IOU values for overlaps.** IOU values are shown for any overlaps between predicted bounding box (blue for CAA, red for Cored) and label bounding box (black). CAA labels are denoted by the "@" symbol, while Cored labels are denoted by the "*" symbol.

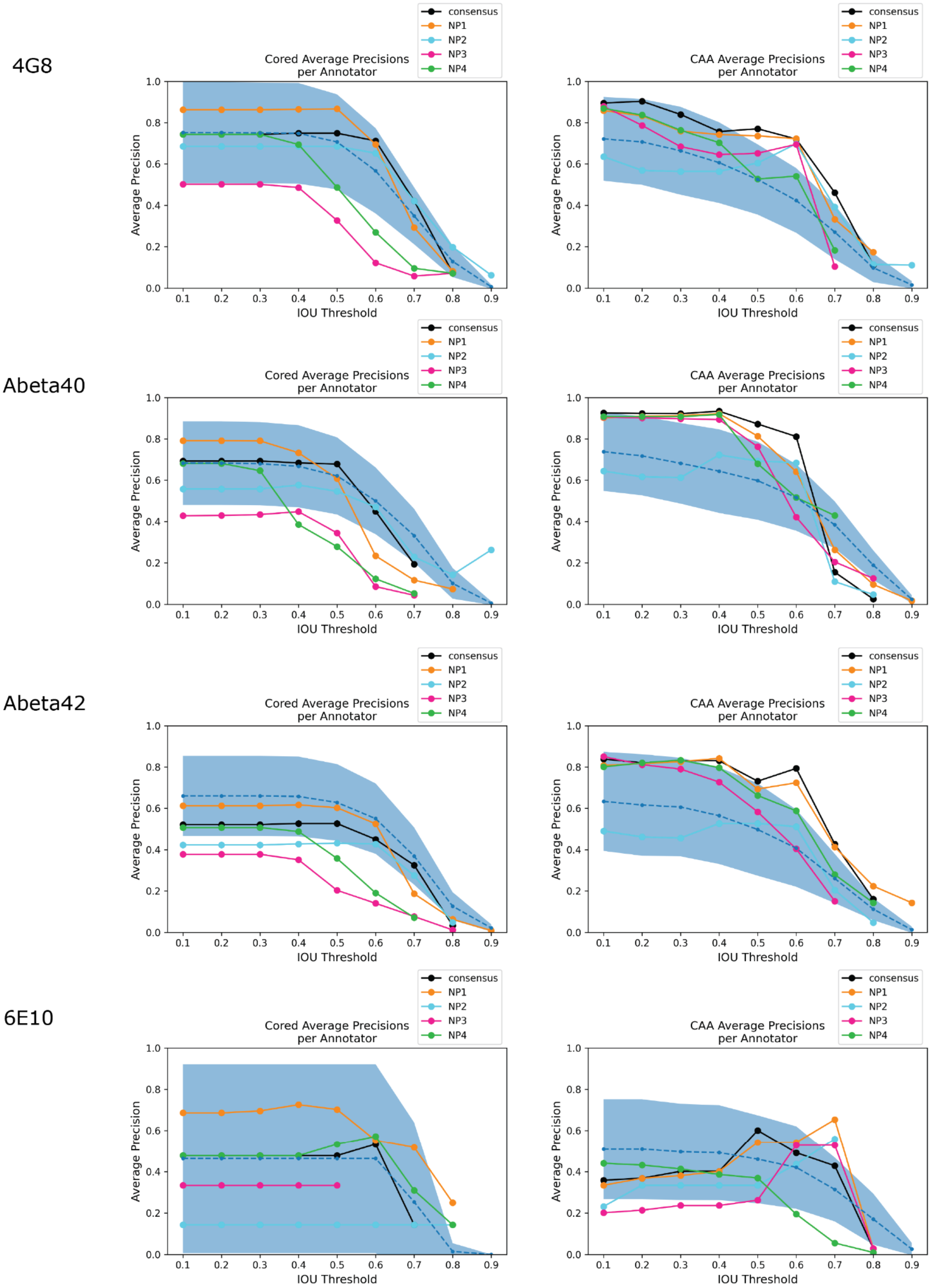


**Supplemental Figure 4: Average precision by stain.** We compute average precision on the prospective validation set stratified by stain. At certain IOU thresholds, there are no true positive cases and correspondingly no precision scores.

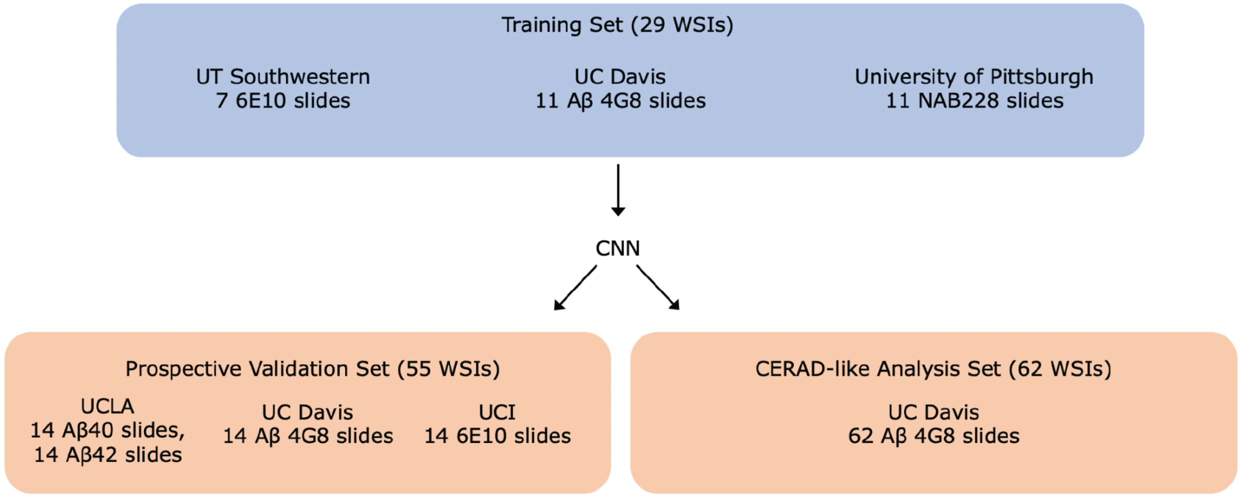


**Supplemental Figure 5: Schematic of training and prospective validation datasets.** We used a total of 29 WSIs from three institutions for training all models from Wong et al[1]. We used a separate new dataset of 55 WSIs from three institutions for prospective validation. To calculate comparison versus CERAD-like scores (Figure 4), we used a third dataset of 62 WSIs from Tang et al[2].

| | |
|---|---|
| Architecture | x86_64 |
| CPU op-mode(s) | 32-bit, 64-bit |
| Byte Order | Little Endian |
| CPUs | 64 |
| Thread(s) per core | 2 |
| Core(s) per socket | 16 |
| Socket(s) | 2 |
| NUMA node(s) | 2 |
| Vendor ID | GenuineIntel |
| CPU family | 6 |
| Model | 79 |
| Model name | Intel(R) Xeon(R) CPU E5-2697A v4 @ 2.60GHz |
| Stepping | 1 |
| CPU MHz | 1200.024 |
| CPU max MHz | 3600.0000 |
| CPU min MHz | 1200.0000 |
| BogoMIPS | 5200.02 |
| Virtualization | VT-x |
| L1d cache | 32K |

**Supplemental Table 1. CPU specifications.**
