## Supplemental File (Protocol) for "Learning fast and fine-grained detection of amyloid neuropathologies from coarse-grained expert labels"

### Amyloid Beta Object Detection

Thank you for volunteering to be an annotator for this deep learning (DL) study that focuses on improving object detection, a critical initial input to developing models that will augment the ability of neuropathology experts!

Your annotations are invaluable, and necessary to validating our DL model which has shown a lot of promise. There are opportunities here to translate advances in the deep learning field to clinical research practice. The algorithm is not only fully interpretable, but also very fast and can even be run on a personal computer (no specialized hardware for deep learning needed!). We hope that this will make technological advances more accessible and equitable for novices and experts alike, and become highly impactful work that will be widely used.

We will be using the platform called "SuperAnnotate" to collect annotations. You should have received an invitation to the project. If not, please. There are 200 total images to annotate. Each image should take about 30 seconds to annotate, for a total time of 1 hour and 40 minutes for the whole project. All images should be annotated by **October 15, 2021.** Annotations can be completed either periodically, or all at once. Please see below for instructions on how to annotate. Thank you!

When you login, the landing page looks like this:

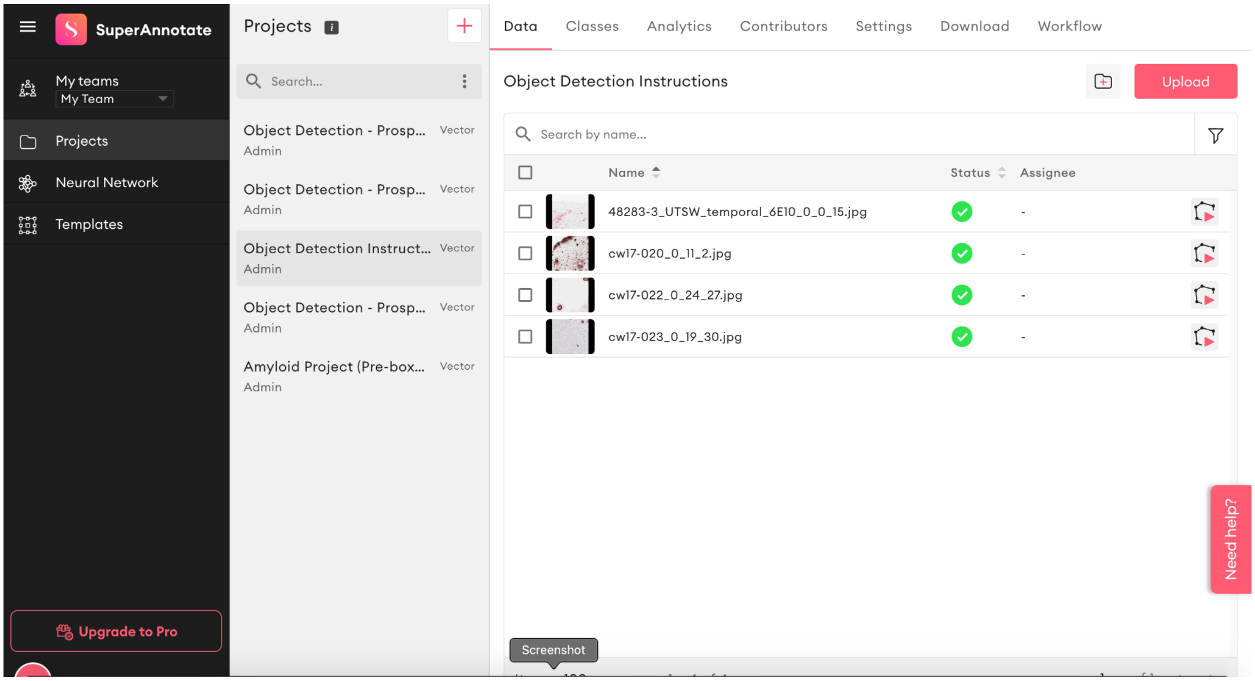


Click on the project entitled “Object Detection - Prospective Validation (YOUR NAME)”. There should only be one project to select from. Select the first image to begin annotating. Once selected, the platform should look something like this:

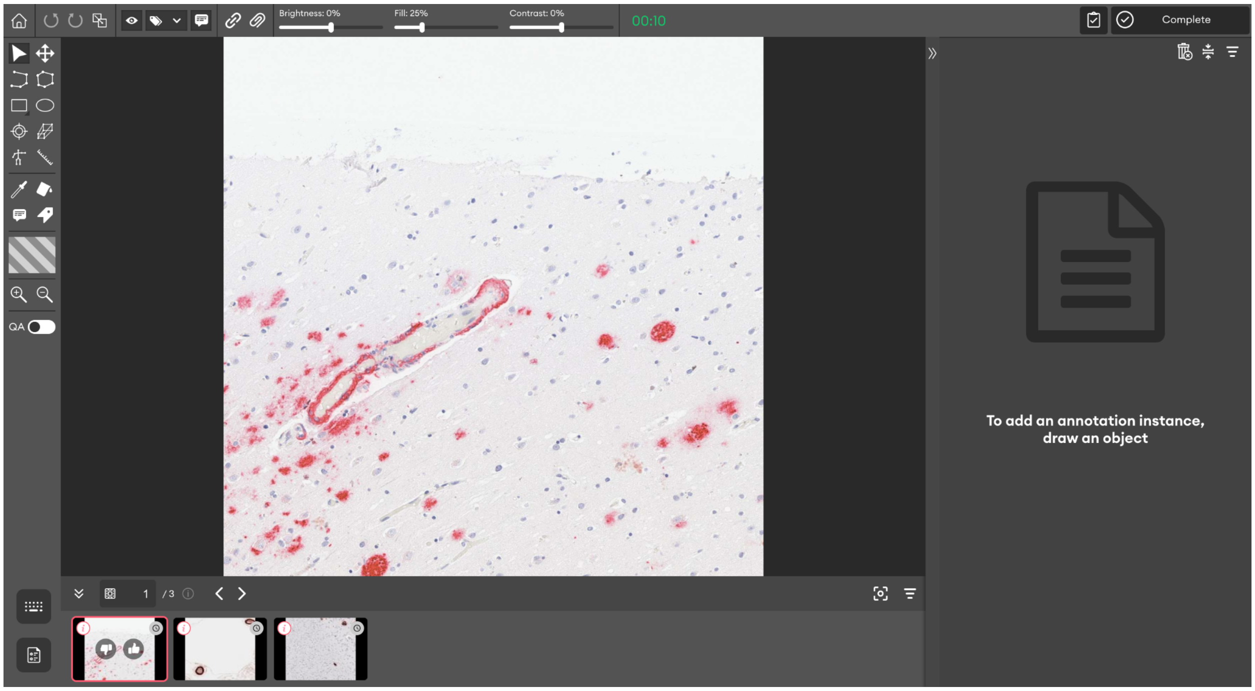

**Instructions:**

1) Identify **ALL Cored Amyloid-β plaques and CAA pathologies** within each image (ignore any other plaque types), and draw an appropriately labeled bounding box around each pathology. Try to capture the entire pathology within the box.

2) To draw a bounding box, you can click this square button indicated with a red arrow in the following image, or simply click the **“X”** key on your keyboard:

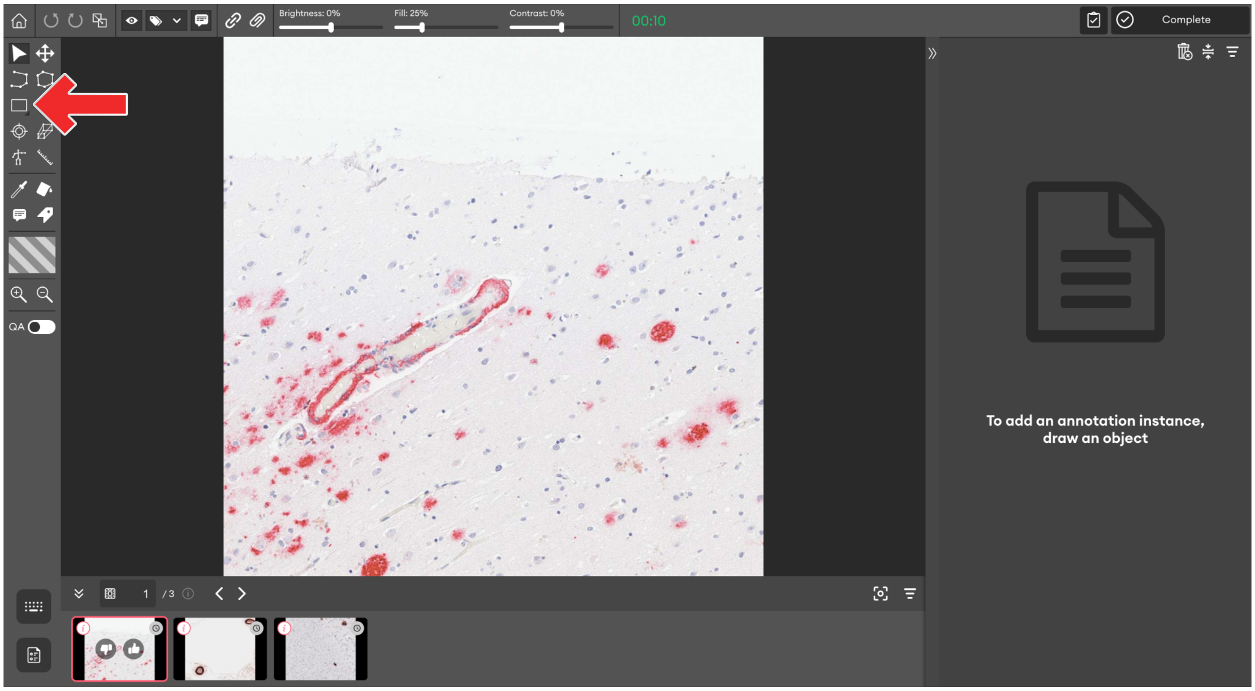


3) Now once the bounding box tool is selected, then simply click and drag to form a box that captures the entire pathology. It should look like this. After dragging to form a box, the box can be shifted by clicking the box and dragging it, or scaled by dragging the corners
    a) If a pathology is only partly within the image and the rest is out of frame, box as much of the pathology as you can.
    b) If there are two pathologies adjacent to each other, use separate boxes - one box for a single pathology.

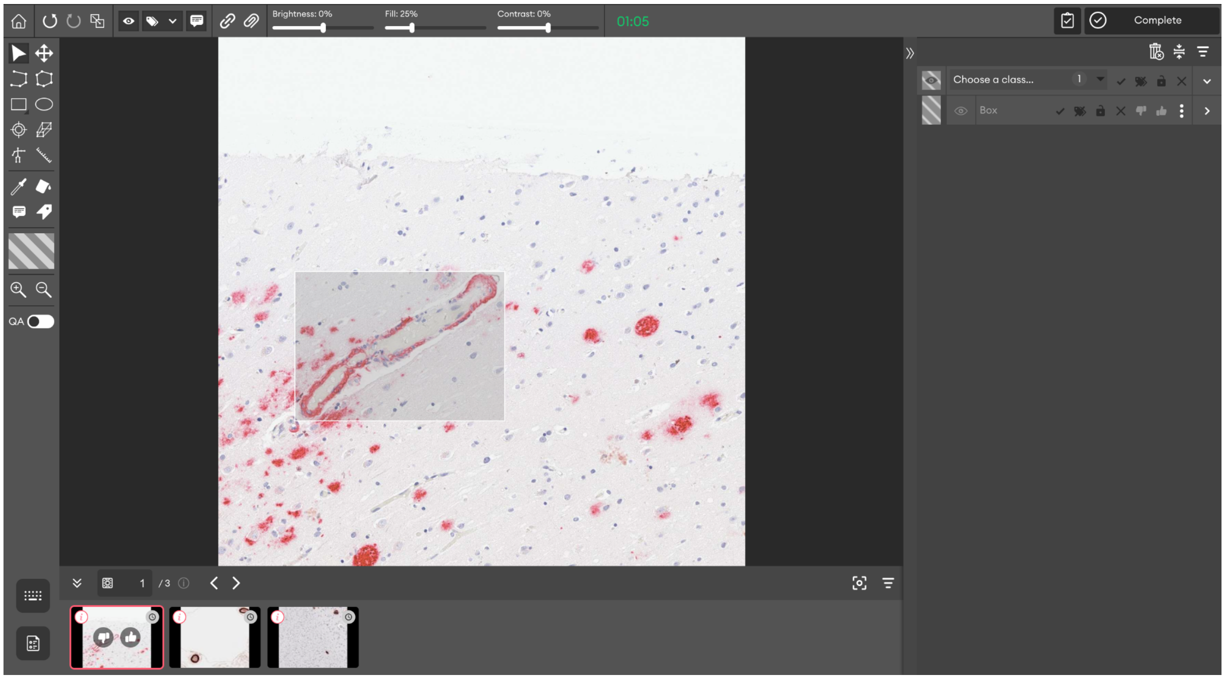


4) Once the box is drawn and captures as much of the pathology as you can, then right click on the box and select which class this pathology belongs to. Alternatively you can set the class by using the right panel and selecting “Choose a class”:

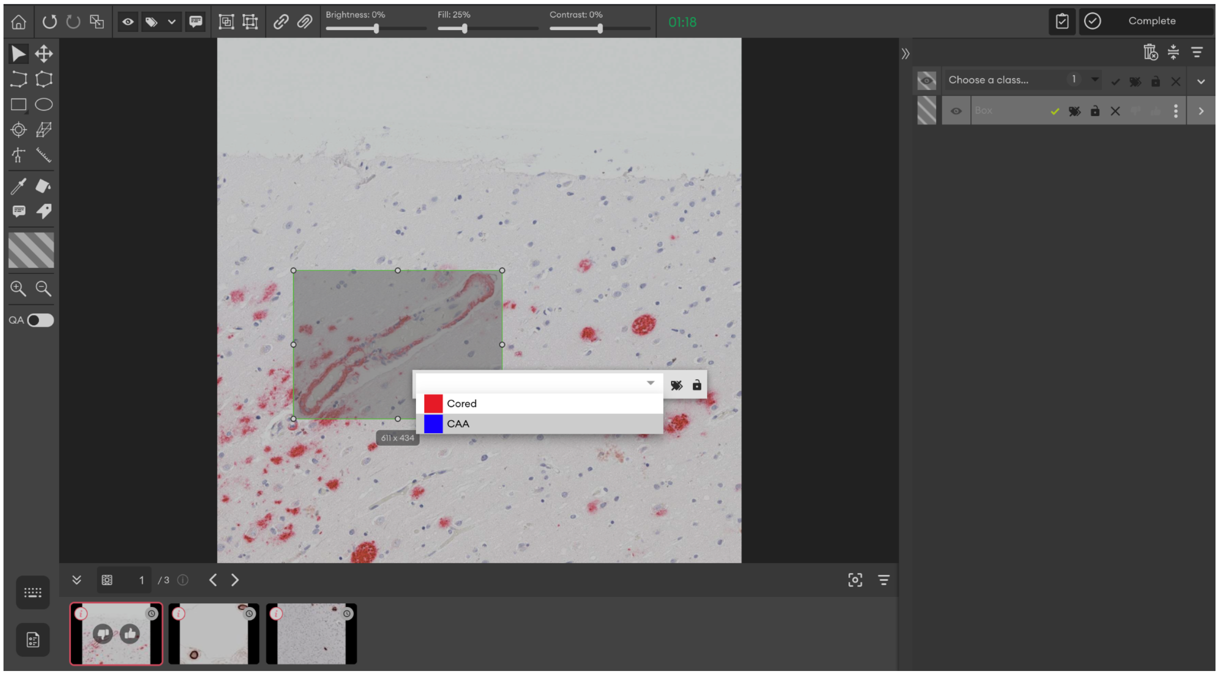


5) Once selected, the portal should look like this:

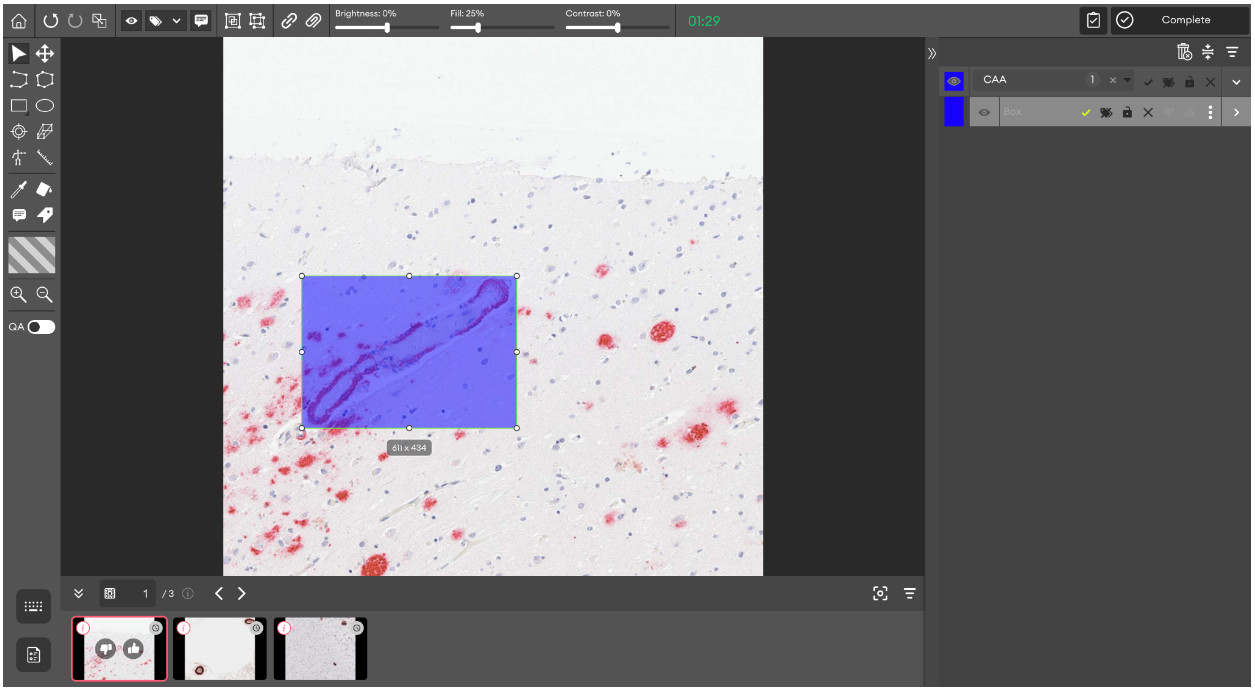


6) Repeat this process for each Cored or CAA pathology in the image.

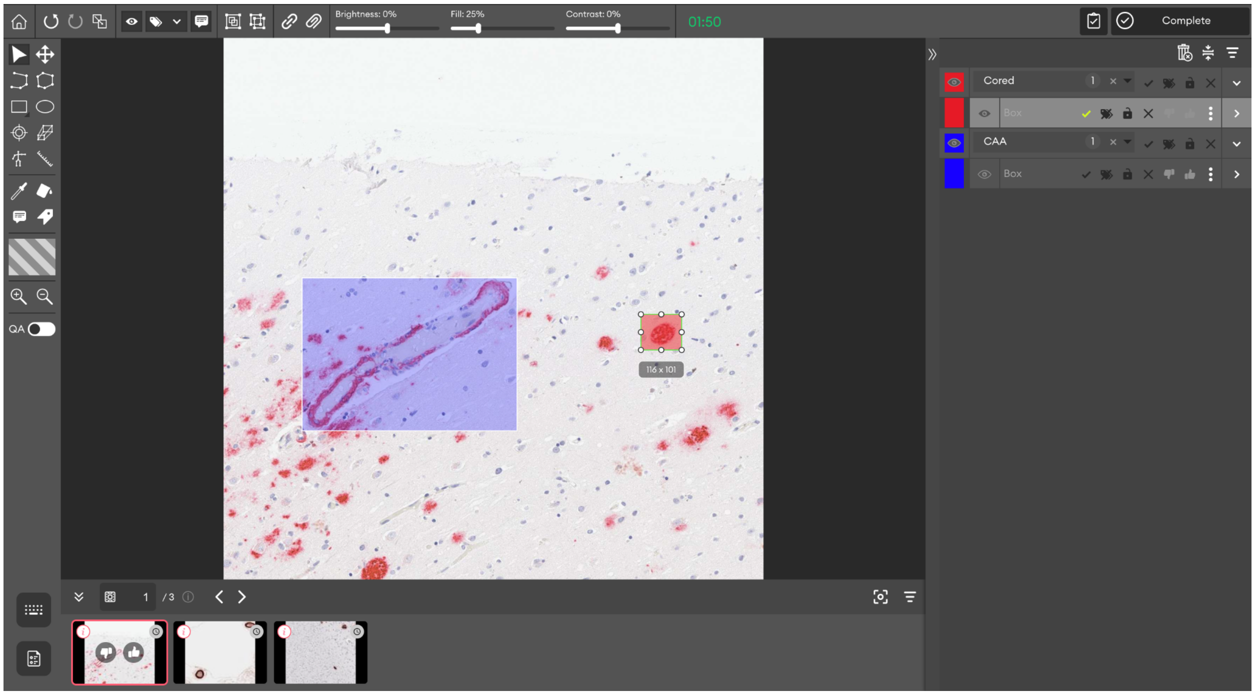

7) Once all pathologies have been captured by a labeled bounding box, move onto the next image.
8) Please make sure that all 200 images are annotated (or if no Cored or CAA pathology exists within an image, then leave the image blank and move onto the next)

**More Examples**

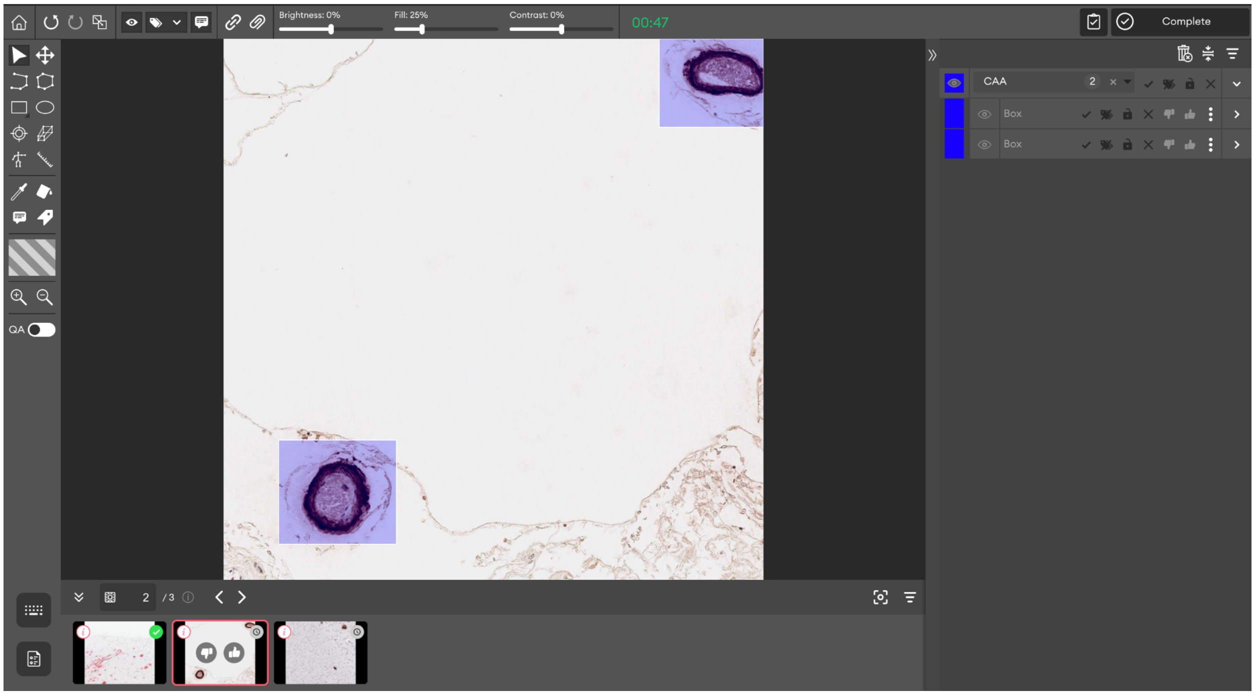


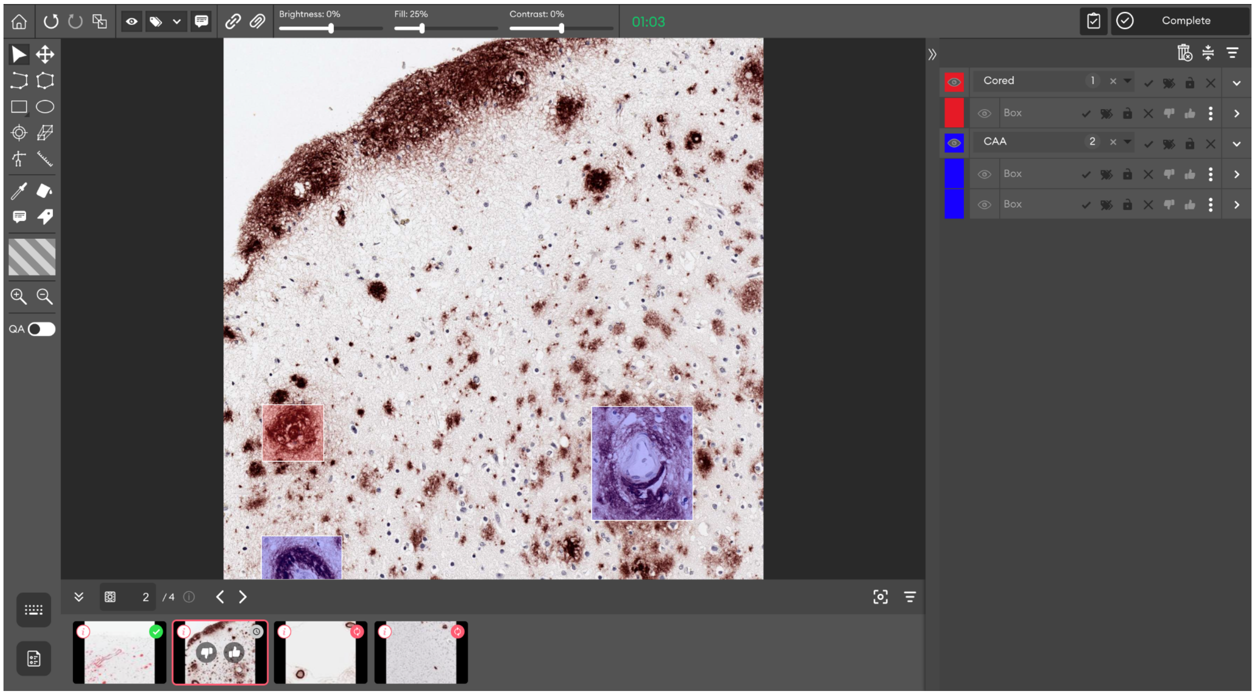

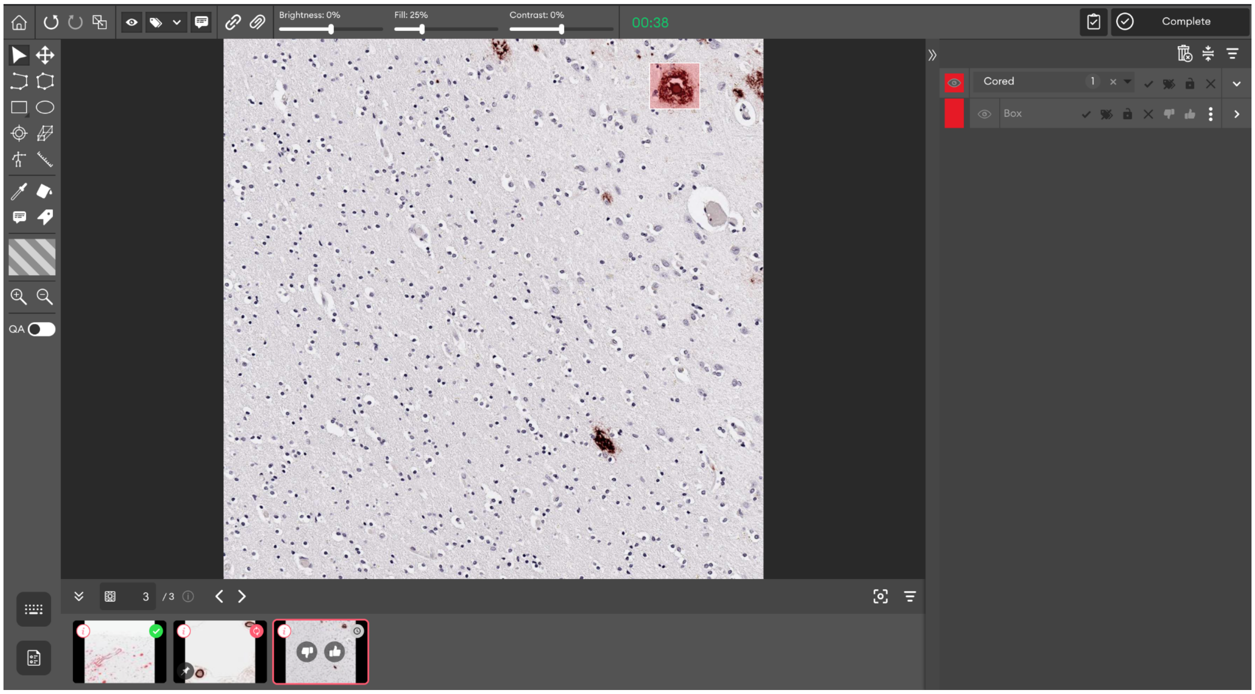


**Other Useful Information:**

- The image can be zoomed using the + and - magnifying glasses on the left tool panel
- The image can be panned using the “drag” button on the left tool panel
- To display all keyboard shortcuts, simply press “Ctrl” + “K” on your keyboard

**Troubleshooting:**

- If you run into any technical issues, try refreshing the page, or clicking the home button on the top left corner of the platform
- If any issues persist, please
